# Concurrent anodal tDCS enhances motor learning and increases cortical TrkB phosphorylation in mice

**DOI:** 10.64898/2026.10.08.757773

**Authors:** Ying Wang, Qingping Yu, Yunqing Wen

## Abstract

The timing of transcranial direct current stimulation (tDCS) relative to motor practice may influence its behavioral effects, but the accompanying molecular responses remain incompletely characterized. Here, we examined cortical current delivery, rotarod performance, and BDNF–TrkB signaling in mice receiving anodal tDCS concurrently with training or during intertrial intervals. Cortical current measurements indicated that approximately half of the applied current reached the measurement site under the experimental stimulation configuration. During four days of rotarod training, anodal tDCS concurrent with motor tasks improved performance at selected trials and increased footstep frequency, whereas stimulation delivered before or after training trials did not significantly improve performance. Preceding online anodal tDCS with cathodal stimulation provided no detectable additional behavioral benefit. Concurrent anodal tDCS increased TrkB phosphorylation in the stimulated cortex after both one and four days, without a significant increase in BDNF levels. Paused offline anodal tDCS stimulation also increased TrkB phosphorylation after one day despite lacking a detectable behavioral benefit. No significant molecular changes were detected in the sampled non-targeted brain regions. Genetic experiments revealed reduced rotarod performance in BDNF heterozygous mice at several trials and trial-dependent responses to online stimulation in BDNF-KIV mice. Together, these findings show that concurrent anodal tDCS improves rotarod performance and increases TrkB phosphorylation in the stimulated cortex. These results motivate further investigation of how BDNF–TrkB signaling interacts with task-related neural activity during stimulation.

**One Sentence Summary:** Concurrent anodal tDCS enhances motor learning in mice, while both concurrent and offline stimulation increase local cortical TrkB phosphorylation.

## Introduction

Transcranial direct current stimulation (tDCS) is a non-invasive neuromodulation technique that delivers weak direct current through electrodes placed on the head. Early studies demonstrated that tDCS can modify human motor cortical excitability, with effects that depend on stimulation polarity, intensity, and duration(*1*). When combined with motor practice, anodal tDCS over the primary motor cortex (M1) can enhance skill acquisition across repeated training sessions, supporting its investigation as an adjunct to motor training and rehabilitation(*2–4*). Understanding how stimulation interacts with ongoing motor practice is therefore essential for interpreting its behavioral effects and developing effective stimulation protocols.

The effects of tDCS involve both acute modulation of neuronal excitability and changes in synaptic plasticity. Direct current can polarize neuronal compartments, including cell bodies, dendrites, and axon terminals, with the magnitude and direction of polarization depending on neuronal morphology and electric field orientation(*5, 6*). Experimental studies further show that subthreshold electric fields can modulate presynaptic release through changes in axonal polarization and calcium-dependent release mechanisms(*7, 8*). In parallel, direct current stimulation can modify activity-dependent synaptic plasticity through interactions with postsynaptic firing and ongoing synaptic activity(*9, 10*). Pharmacological studies in humans have further implicated N-methyl-D-aspartate receptors (NMDARs) in the excitability changes that persist after stimulation (*11*). These findings suggest that the effects of tDCS cannot be explained solely by a uniform increase or decrease in cortical excitability, but also depend on how electrical stimulation interacts with cellular and synaptic activity.

This interaction provides a rationale for delivering stimulation concurrently with task performance. In our previous study, concurrent anodal tDCS enhanced motor learning in a task-specific manner and preferentially increased the activity of task-related M1 neurons during movement (*12*). These findings suggested that stimulation can facilitate plasticity within circuits engaged by the ongoing task. However, the molecular responses accompanying this behavioral enhancement, and their relationship to stimulation timing, remain incompletely characterized.

Brain-derived neurotrophic factor (BDNF) and its receptor, tropomyosin receptor kinase B (TrkB), are candidate components of this activity-dependent response. In mouse motor cortical slices, direct current stimulation paired with repeated synaptic activation induces potentiation that requires NMDAR activation, activity-dependent BDNF secretion, and TrkB activation (*13*). In a separate mouse study, anodal tDCS enhanced hippocampal synaptic plasticity and memory alongside epigenetic regulation of BDNF expression(*14*). Other cellular mechanisms may also contribute: tDCS-induced enhancement of sensory cortical responses has been linked to astrocytic calcium signaling(*15*). Together, these studies provide a basis for investigating BDNF – TrkB signaling during motor training, while indicating that stimulation can engage multiple cellular processes.

Although concurrent stimulation can enhance motor learning and task-related cortical activity(*12*), it remains unclear whether the accompanying molecular responses differ from those elicited by stimulation delivered outside motor practice. Activity-dependent BDNF release has been demonstrated in neuronal preparations, and BDNF secretion and TrkB activation have been implicated in direct current stimulation-associated synaptic potentiation(*13, 16*). However, BDNF abundance in tissue homogenates does not directly measure its release, and BDNF abundance and TrkB phosphorylation capture distinct aspects of the pathway. Comparing these measures across stimulation schedules and brain regions may therefore help clarify the relationship between cortical signaling and behavioral improvement. In addition, examining mice with genetic alterations affecting BDNF or TrkB can characterize how changes in this system relate to baseline task performance and responses to stimulation.

Here, we combined cortical current measurements, behavioral analysis, and molecular assays to investigate anodal tDCS during rotarod training in mice. We first characterized the current reaching the cortical measurement site under our electrode configuration. We then compared stimulation delivered concurrently with training with stimulation delivered during intertrial intervals and quantified stepping behavior alongside rotarod performance. To examine accompanying molecular responses, we measured TrkB phosphorylation and BDNF levels in the stimulated cortex and additional brain regions after one or four days of stimulation. Finally, we assessed rotarod performance and stimulation responses in BDNF heterozygous, TrkB heterozygous, and BDNF-KIV mice. These experiments were designed to examine whether the behavioral effects of anodal tDCS depend on its timing relative to motor practice, how these effects relate to cortical BDNF–TrkB signaling, and how genetic alterations affecting this system influence task performance and responses to stimulation.

## Materials and Methods

### Animals

Male wild-type C57BL/6J mice aged 7–10 weeks were obtained from Shanghai SLAC Laboratory Animal Co., Ltd. (Shanghai, China). Mice were randomly assigned to active stimulation or the corresponding sham groups. Separate cohorts were used for different experimental protocols and tissue collection time points. Male BDNF heterozygous (Bdnf+/−), TrkB heterozygous (Ntrk2+/−), and BDNF-KIV mice were obtained from existing breeding colonies maintained in our laboratory. All mice used in the genetic experiments were 7–10 weeks old and maintained on a C57BL/6 background. BDNF and TrkB heterozygous mice were compared with their corresponding wild-type littermates (Bdnf+/+ and Ntrk2+/+, respectively), and BDNF-KIV mice were compared with wild-type controls. Mice were housed under a 12-h light/dark cycle at 19–22 °C, with lights on from 07:00 to 19:00. Behavioral experiments were conducted during a consistent time window each day. All animal procedures were approved by the Animal Committee of the Institute of Neuroscience, Center for Excellence in Brain Science and Intelligence Technology, Chinese Academy of Sciences. Procedures were performed in accordance with institutional guidelines, with efforts to minimize animal use and discomfort.

### Electrode implantation

The electrode configuration and rotarod training procedures followed our previously published protocol, with the modifications described below(*12*). Mice were anesthetized with sodium pentobarbital administered intraperitoneally at 7 mg/kg. Animals were positioned in a stereotaxic frame (model 68030, RWD Life Science Co., Ltd., Guangdong, China). Body temperature was maintained using a heating pad at 38 °C. The scalp and underlying connective tissue were removed to expose the skull. A tubular plastic electrode holder with an internal contact area of 3.14 mm² was secured over the right M1. The center of the holder was positioned at the anteroposterior level of bregma and 1.5 mm lateral to the midline. A round tin return electrode was implanted subcutaneously on the contralateral side of the back of the neck. Its dimensions were approximately 5 mm in diameter. Mice were allowed to recover for seven days before training and stimulation.

### Transcranial direct current stimulation

Before stimulation, the epicranial electrode holder was filled with 0.9% NaCl solution. tDCS was delivered using a constant-current stimulator (model ST1, Quanlan Technology Co., Ltd., Shanghai, China). For anodal stimulation, the electrode over M1 served as the anode, and the subcutaneous return electrode served as the cathode. Electrode polarity was reversed for cathodal stimulation. Unless otherwise indicated, stimulation was delivered at 0.1 mA. With an epicranial contact area of 3.14 mm^2^, this corresponded to a nominal electrode current density of approximately 3.2 mA/cm^2^.

For online stimulation, anodal tDCS was initiated at the beginning of the familiarization session and each rotarod training trial. Each stimulation episode lasted 5 min, regardless of whether the mouse fell from the rod before the trial reached its maximum duration, ensuring a fixed stimulation duration across animals. Thus, stimulation continued after an early trial termination. No additional stimulation was delivered between these scheduled episodes.

For pre-trial offline stimulation, anodal tDCS was delivered for 5 min before familiarization and during each 5-min interval preceding a training trial. For post-trial offline stimulation, anodal tDCS was delivered for 5 min following familiarization and each training trial. No current was delivered during task performance in either offline protocol.

For the combined cathodal–anodal protocol, mice received 5 min of cathodal tDCS before familiarization and before each training trial, followed by 5 min of anodal tDCS initiated at the start of familiarization or the training trial. Both polarities were delivered at 0.1 mA. Each anodal stimulation episode was completed regardless of early trial termination. Current was switched on and off directly, without ramp-up or ramp-down periods. Sham mice underwent the same electrode connection, handling, and training procedures without current delivery. Animals were monitored for abnormal behavior and signs of discomfort throughout stimulation.

### Accelerating rotarod training

A five-lane accelerating rotarod with a rod diameter of 3 cm (model 47600, Ugo Basile, Gemonio, Italy) was used to assess motor performance. Mice were habituated to the testing room for 2 h before testing. Each daily session began with 5 min of familiarization at a constant speed of 4 rpm on days 1–2 and 8 rpm on days 3–4. Training consisted of three trials per day for four consecutive days, separated by 5-min rest intervals off the rod. The rotarod accelerated linearly from 4 to 40 rpm on days 1–2 and from 8 to 80 rpm on days 3–4 during each 300-s trial. Each trial ended when the mouse fell from the rod or reached the maximum trial duration of 300 s. Time spent on the rod was recorded for each trial. To assess the persistence of behavioral benefits after stimulation, mice were tested on day 18, 14 days after completion of the four-day training and stimulation protocol. The follow-up session consisted of three accelerating rotarod trials at 8–80 rpm, separated by 5-min intertrial intervals. No tDCS was delivered during the follow-up session.

### Video analysis of stepping behavior

Video analysis was performed on trials 1, 6, 7, and 12 of the four-day rotarod training protocol, which consisted of three trials per day. Trials 1 and 6 corresponded to the first and last trials of training at 4–40 rpm (days 1–2), respectively, whereas trials 7 and 12 corresponded to the first and last trials of training at 8–80 rpm (days 3–4). For each selected trial, the first 80 s were analyzed, as all mice remained on the rod throughout this period. This fixed observation window ensured that measurements were obtained over the same elapsed time and corresponding rotarod speed range within each selected trial. Only the left hindpaw, contralateral to the stimulated right M1, was analyzed. Step timing and footstep frequency were calculated separately for each mouse at each selected trial. Individual steps were summarized within each mouse before group comparisons, with the mouse serving as the experimental unit.

### Measurement of current reaching the cortex

Current beneath the epicranial stimulation site was measured in awake mice using the electrode configuration illustrated in Fig. 1. During electrode implantation, a circular skull flap approximately 2.5 mm in diameter was temporarily lifted over the right primary motor cortex (M1). A custom-built recording electrode corresponding to the smallest electrode design shown in Fig. 1 was inserted beneath the skull flap and positioned over the intact dura mater. The skull flap was then replaced, leaving the recording electrode between the skull and dura, directly beneath the epicranial stimulation site. The epicranial stimulation contact was 2 mm in diameter, and 0.9% NaCl solution was used as the contact medium.

**Fig. 1.**
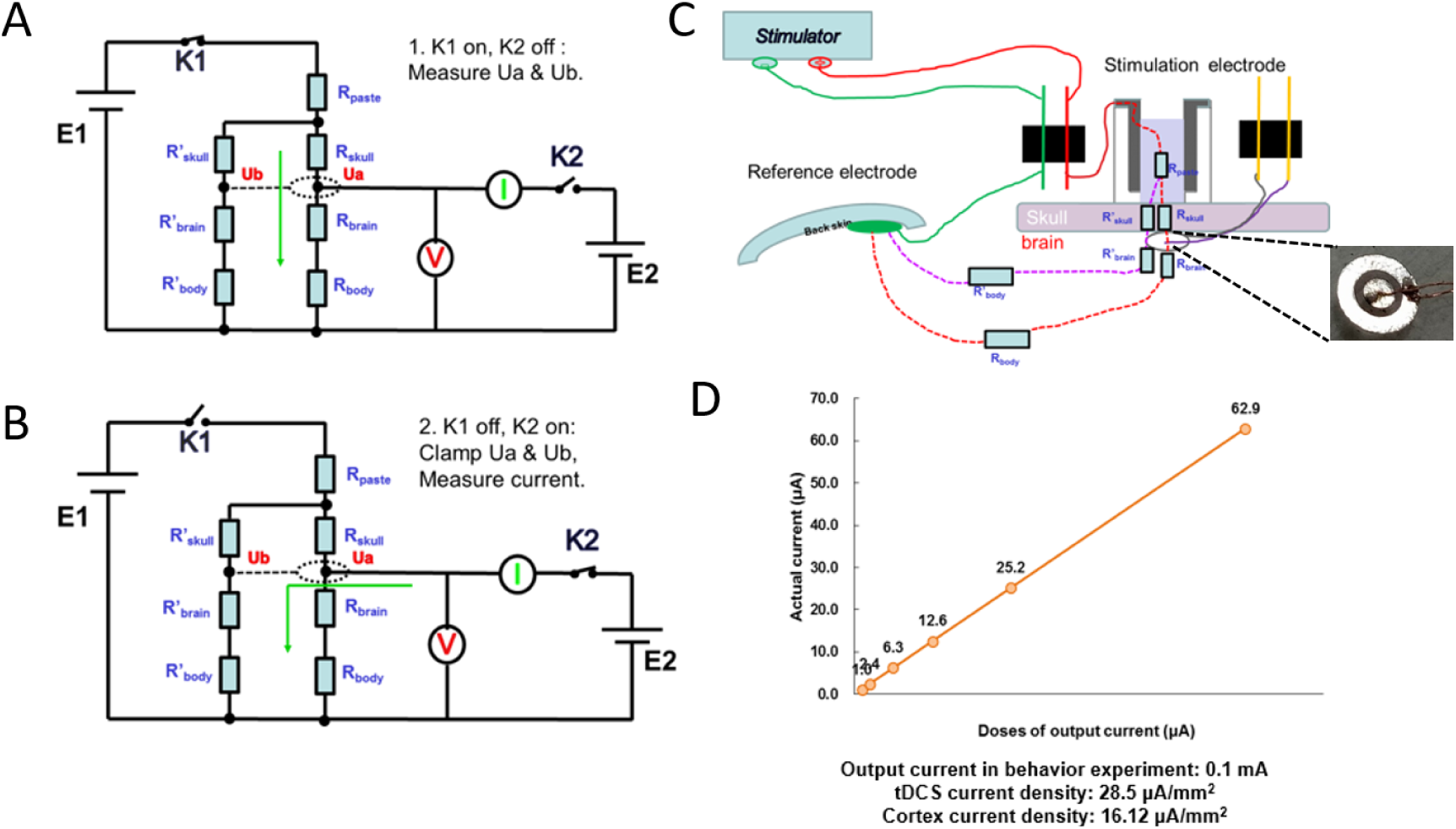
Experimental configuration and measurement of cortical current during transcranial direct current stimulation in mice. **(A, B)** Equivalent circuit diagrams illustrating the two-step measurement procedure. **(A)** With switch K1 closed and K2 open, potentials Ua and Ub were measured during stimulation. **(B)** With K1 open and K2 closed, the previously measured potentials were reproduced by voltage clamping, and the resulting current was recorded. Rpaste denotes the resistance of the conductive paste; Rskull, Rbrain, and Rbody, together with their primed counterparts, represent the tissue resistances in the two modeled current pathways. E1 and E2 denote the voltage sources, and V and I indicate voltage and current measurements, respectively. **(C)** Schematic of the stimulation and measurement configuration, showing the electrode arrangement and modeled current pathways through the skull, brain, and body. The inset shows the electrode assembly. **(D)** Measured current as a function of stimulator output current. Open circles represent measurements at the tested output levels, with adjacent points connected by a line.

The stimulation electrode montage matched that used in the behavioral experiments: the anode was positioned over the right M1, and the return electrode was placed subcutaneously over the contralateral back. Measurements were obtained using a custom-built current-measurement system operated with an in-house program written in LabVIEW (National Instruments). Applied stimulation currents were 1.0, 2.4, 6.3, 12.6, and 25.2 μA, with each stimulation lasting 10 s.

The recorded-to-applied current ratio was calculated by dividing the current measured beneath the stimulation site by the applied stimulation current and expressing the result as a percentage. Measurements were obtained from two mice, with six repeated measurements at each applied current amplitude in each mouse.

An additional measurement was performed at an applied stimulation current of 0.1 mA, matching the current used in the behavioral experiments. The estimated cortical current density beneath the stimulation site was 16.12 μA/mm². [Verify the conversion from measured current to current density, including the effective recording area.] The cortical-to-epicranial current-density ratio was calculated by dividing the estimated cortical current density by the current density at the epicranial stimulation contact.

### Tissue collection

For the molecular experiments shown in Fig. 3 and Supplementary Fig. 2, tissue collection procedures were initiated immediately after completion of the final behavioral session following one or four days of training and stimulation. Mice were anesthetized with sodium pentobarbital administered intraperitoneally at 7 mg/kg, and brains were removed without perfusion. Tissue samples were dissected from the stimulated right primary motor cortex (M1), the corresponding left M1, and the ipsilateral and contralateral distal cortical regions indicated in Fig. 3A. For the regional comparison shown in Fig. 3F, tissue was collected from the right M1, left M1, and auditory cortex in both hemispheres. For the molecular analysis shown in Fig. 4D, tissue collection followed the same procedure after the final training and stimulation session on day 4. Samples were obtained from the targeted right M1. In the stimulated BDNF-KIV group, anodal tDCS was delivered according to the online stimulation protocol described above. All dissected tissue samples were stored at −80 °C until processing for western blot analysis.

### Western blotting

Brain tissue samples were homogenized on ice in RIPA lysis buffer supplemented with protease and phosphatase inhibitors. Homogenates were centrifuged at 12,000 × g for 15 min at 4 °C, and the supernatants were collected. Protein concentrations were determined using a bicinchoninic acid (BCA) assay. Equal amounts of protein (20 μg per lane) were mixed with SDS–PAGE sample buffer and heated at 95 ℃ for 5 min. Proteins were separated by SDS–PAGE, using 8% gels for phosphorylated and total TrkB and 12–15% gels for BDNF and actin, and transferred to PVDF membranes. Membranes were blocked for 1 h at room temperature in Tris-buffered saline containing 0.1% Tween-20 (TBST) and 5% bovine serum albumin for phosphoprotein detection or 5% non-fat milk for other targets. Membranes were incubated overnight at 4 ℃ with primary antibodies against phosphorylated TrkB, total TrkB, BDNF, and actin. After three washes with TBST, membranes were incubated with appropriate horseradish peroxidase-conjugated secondary antibodies for 1 h at room temperature and washed again with TBST. Immunoreactive bands were visualized using enhanced chemiluminescence and recorded with a chemiluminescence imaging system.

Band intensities were quantified using ImageJ following background subtraction. Phosphorylated TrkB signals were normalized to the corresponding total TrkB signals, and BDNF signals were normalized to actin. Normalized values were expressed relative to the corresponding control group within each experiment. Each mouse was treated as an independent biological replicate for statistical analysis.

### Statistical analysis

Data are presented as mean ± standard error of the mean (SEM). Independent-samples *t*-tests were used for the between-group comparisons indicated in the figures. Tests were two-tailed, and Student’s *t*-tests were used. The mouse was the biological experimental unit. Repeated stepping measurements and technical measurements were summarized within each mouse before group comparisons. Sample sizes are reported in the corresponding figures. Statistical significance was defined as P < 0.05. Analyses were performed using GraphPad Prism (Version 5.0, GraphPad, San Diego, CA, USA].

## Results

### Approximately half of the applied current reaches the cortical measurement site

To characterize current delivery under the stimulation configuration used in this study, we measured the current reaching the cortex during stimulation applied over the skull. The measured cortical current was approximately 50% of the applied current (Fig. 1). These measurements characterized current transmission under the experimental electrode configuration and provided a physical basis for the subsequent behavioral and molecular experiments.

### Online anodal tDCS improves rotarod performance and increases footstep frequency

We first examined the behavioral effects of anodal tDCS delivered concurrently with rotarod training. Mice underwent three trials per day for four consecutive days, with the rotarod accelerating from 4 to 40 rpm on days 1–2 and from 8 to 80 rpm on days 3–4. Stimulation was delivered during familiarization and task performance (Fig. 2A).

**Fig. 2.**
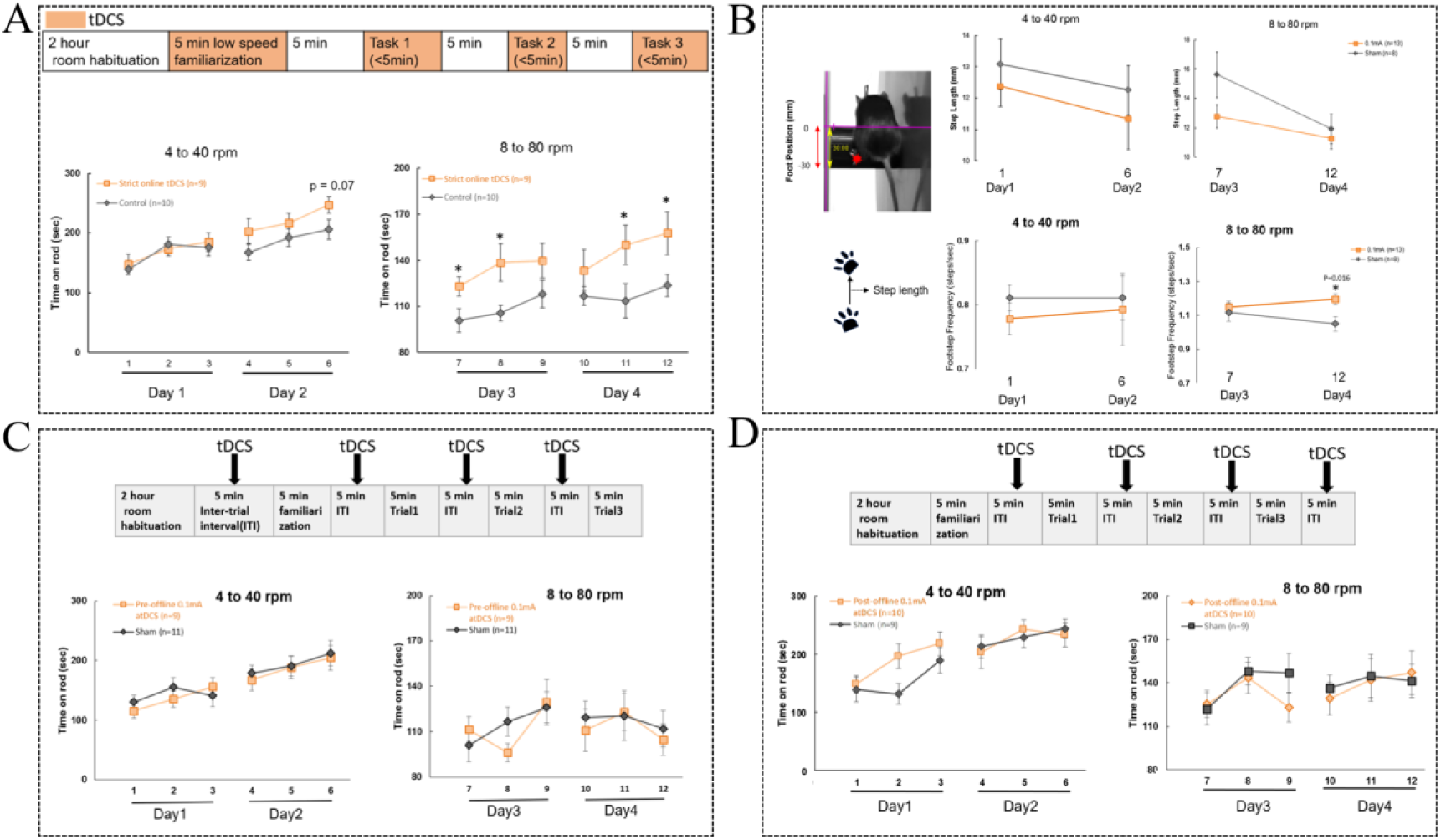
Online anodal tDCS improves rotarod performance and increases footstep frequency. **(A)** Experimental schedule and rotarod performance under the online anodal tDCS protocol. Following 2 h of room habituation, mice underwent 5 min of low-speed familiarization and three training trials, each lasting up to 5 min and separated by 5-min intertrial intervals (ITIs). Orange shading indicates the application of anodal tDCS (0.1 mA) during familiarization and task performance. Mice completed three trials per day for four consecutive days, with the rotarod accelerating from 4 to 40 rpm on days 1–2 and from 8 to 80 rpm on days 3–4. Time on the rod is shown for the online tDCS group (n = 9) and control group (n = 10). **(B)** Video-based analysis of stepping characteristics during rotarod running. The representative image and schematic illustrate the measurements of step length, and step frequency. Plots show step length, and footstep frequency at the indicated trials in mice receiving online anodal tDCS or sham stimulation. Sample sizes are indicated in the plots. **(C)** Experimental schedule and rotarod performance under the pre-task offline protocol. Anodal tDCS (0.1 mA) was applied during 5-min intervals preceding familiarization and each training trial, without stimulation during task performance. Time on the rod is shown for the pre-task offline tDCS group (n = 9) and sham group (n = 11). **(D)** Experimental schedule and rotarod performance under the post-task offline protocol. Anodal tDCS (0.1 mA) was applied during 5-min intervals following familiarization and each training trial. Time on the rod is shown for the post-task offline tDCS group (n = 10) and sham group (n = 9). Data are presented as mean ± SEM. Between-group comparisons at each indicated trial or time point were performed using independent-samples t-tests. *P < 0.05 versus the corresponding control or sham group. Exact P values are provided where indicated. n denotes the number of mice.

At 4–40 rpm, the online tDCS and control groups showed broadly similar performance during the initial trials. A difference favoring online tDCS emerged by the final trial on day 2, although it did not reach statistical significance (P = 0.07). At 8–80 rpm, mice receiving online tDCS remained on the rod significantly longer than controls at several trials on days 3–4 (Fig. 2A).

Video-based analysis was used to examine stepping behavior during the task. Footstep frequency was significantly higher in the online tDCS group on day 4 (P = 0.016), whereas no significant between-group differences were detected in step length or average foot position at the sampled time points (Fig. 2B). Thus, improved rotarod performance was accompanied by an increase in footstep frequency.

We next examined stimulation delivered outside task performance. Anodal tDCS administered during the intervals preceding or following training trials did not significantly improve time on the rod relative to the corresponding sham groups (Fig. 2C, D). This contrasted with the improvements observed during online stimulation.

We also tested whether 5 min of cathodal stimulation before familiarization and each training trial modified the effects of online anodal tDCS (Fig. S1A). Both online anodal tDCS alone and the combined cathodal – anodal protocol produced higher performance than sham stimulation at selected trials. No significant additional improvement was detected with preceding cathodal stimulation compared with online anodal tDCS alone (Fig. S1B). At the day-18 follow-up, conducted without stimulation, mice that had received online anodal tDCS alone or preceding cathodal stimulation followed by online anodal tDCS during training remained on the rod significantly longer than sham-treated mice at selected trials (Fig. S1B). These findings indicate that the behavioral advantage remained detectable two weeks after completion of training and stimulation.

### Anodal tDCS increases TrkB phosphorylation in the stimulated cortex

To examine molecular responses associated with stimulation, we measured TrkB phosphorylation and BDNF levels in the stimulated cortex and additional brain regions after one or four days of training and stimulation (Fig. 3A). TrkB phosphorylation was assessed as the ratio of phosphorylated TrkB to total TrkB, and BDNF signals were normalized to actin.

**Fig. 3.**
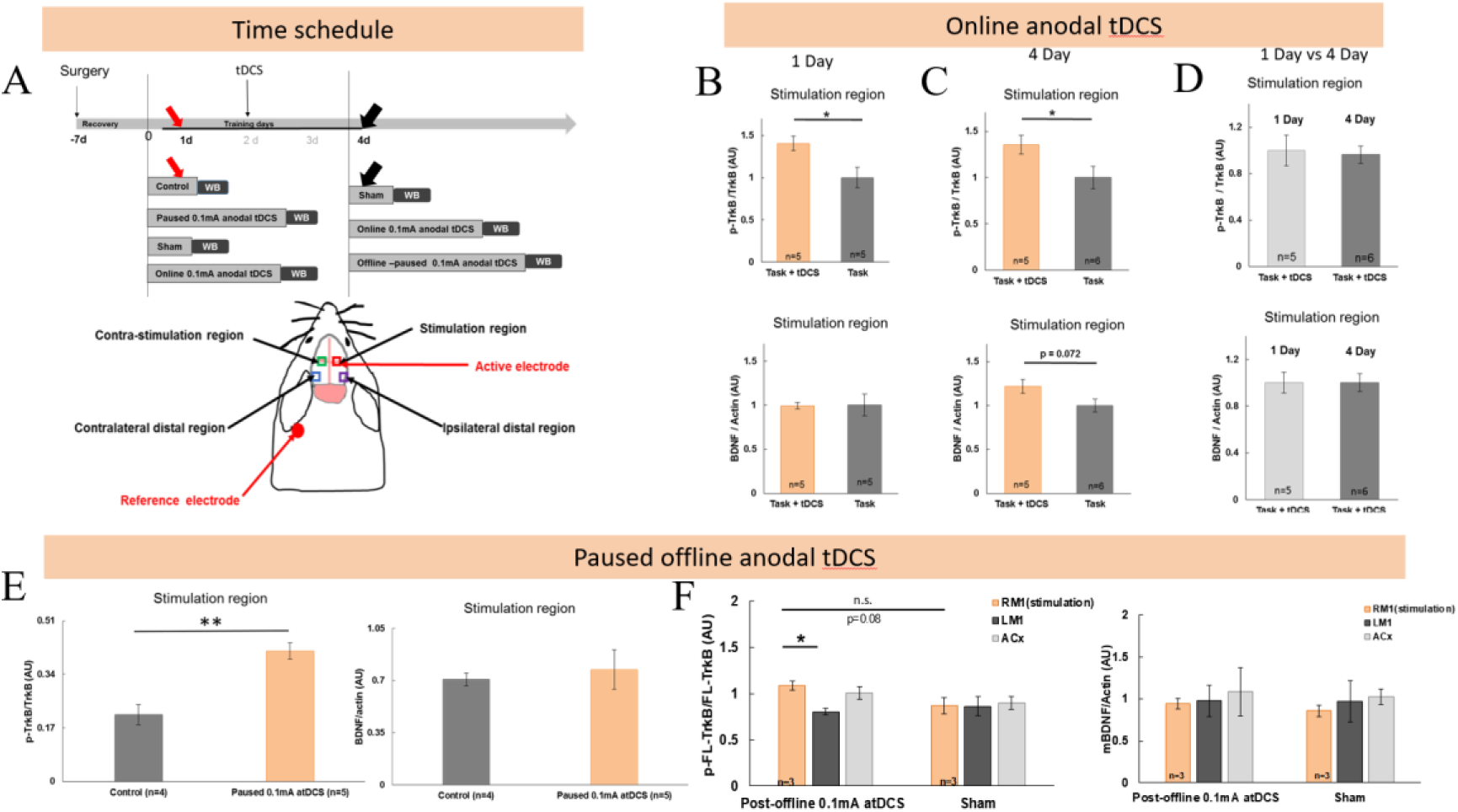
Anodal tDCS increases TrikB phosphorylation in the stimulated cortex. **(A)** Experimental timeline and schematic of electrode placement and cortical tissue sampling. Following electrode implantation and a 7-day recovery period, mice underwent the indicated stimulation and training protocols. Cortical tissue was collected for Western blot analysis after one or four days. The sampling schematic identifies the stimulated region, the corresponding contralateral region, and ipsilateral and contralateral distal regions. **(B, C)** Quantification of TrkB phosphorylation and BDNF abundance in the stimulated region after one day (B) or four days (C) of online anodal tDCS (0.1 mA) during rotarod training. TrkB phosphorylation is expressed as the ratio of phosphorylated TrkB to total TrkB, and BDNF abundance as the ratio of BDNF to actin. The task + tDCS group is compared with the task-only sham group. **(D)** Comparison of TrkB phosphorylation and BDNF abundance in the stimulated region between the one-day and four-day online tDCS groups. **(E)** Quantification of TrkB phosphorylation and BDNF abundance in the stimulated region after one day of paused offline anodal tDCS (0.1 mA), compared with the control group. **(F)** Regional quantification of TrkB phosphorylation and BDNF abundance after four days of post-task offline anodal tDCS (0.1 mA) or sham stimulation. Phosphorylation of full-length TrkB is expressed as the ratio of phosphorylated full-length TrkB to total full-length TrkB. RM1, right primary motor cortex (stimulated region); LM1, left primary motor cortex; ACx, auditory cortex. Values are expressed in arbitrary units (AU). Data are presented as mean ± SEM. Sample sizes are indicated in the plots, and n denotes the number of mice. Comparisons were performed using independent-samples t-tests. *P < 0.05; **P < 0.01; n.s., not significant. Brackets indicate the comparisons tested, and exact P values are shown where indicated.

Online anodal tDCS significantly increased the p-TrkB/TrkB ratio in the stimulated cortex after one day compared with task performance alone (Fig. 3B). An increase was also detected after four days of online stimulation (Fig. 3C). BDNF levels did not differ significantly between groups after one day. After four days, BDNF levels were numerically higher in the online tDCS group, but the difference did not reach statistical significance (P = 0.072). Measurements from the one- and four-day online stimulation groups are presented in Fig. 3D.

Paused anodal tDCS also significantly increased TrkB phosphorylation in the stimulated cortex after one day, without a significant change in BDNF levels (Fig. 3E). Following four days of post-trial offline stimulation, the p-TrkB/TrkB ratio was higher in the stimulated right M1 than in the contralateral left M1. Regional BDNF measurements are shown alongside these comparisons (Fig. 3F).

In the additional sampled regions, no significant differences in TrkB phosphorylation or BDNF levels were detected between paused stimulation and control, between online stimulation and sham, or between online and paused stimulation (Fig. S2). These findings identify a molecular response in the stimulated cortex, while showing that increased TrkB phosphorylation can also occur under a paused stimulation protocol that does not significantly improve rotarod performance.

### Altered BDNF–TrkB signaling is associated with differences in rotarod performance and stimulation responses

We next examined rotarod performance in mice with genetic alterations affecting the BDNF – TrkB system. Without stimulation, BDNF heterozygous mice showed lower performance than wild-type mice, with significant differences at several training trials. In the cohorts receiving concurrent anodal tDCS, no significant genotype differences were detected at the trials shown (Fig. 4A).

**Fig. 4.**
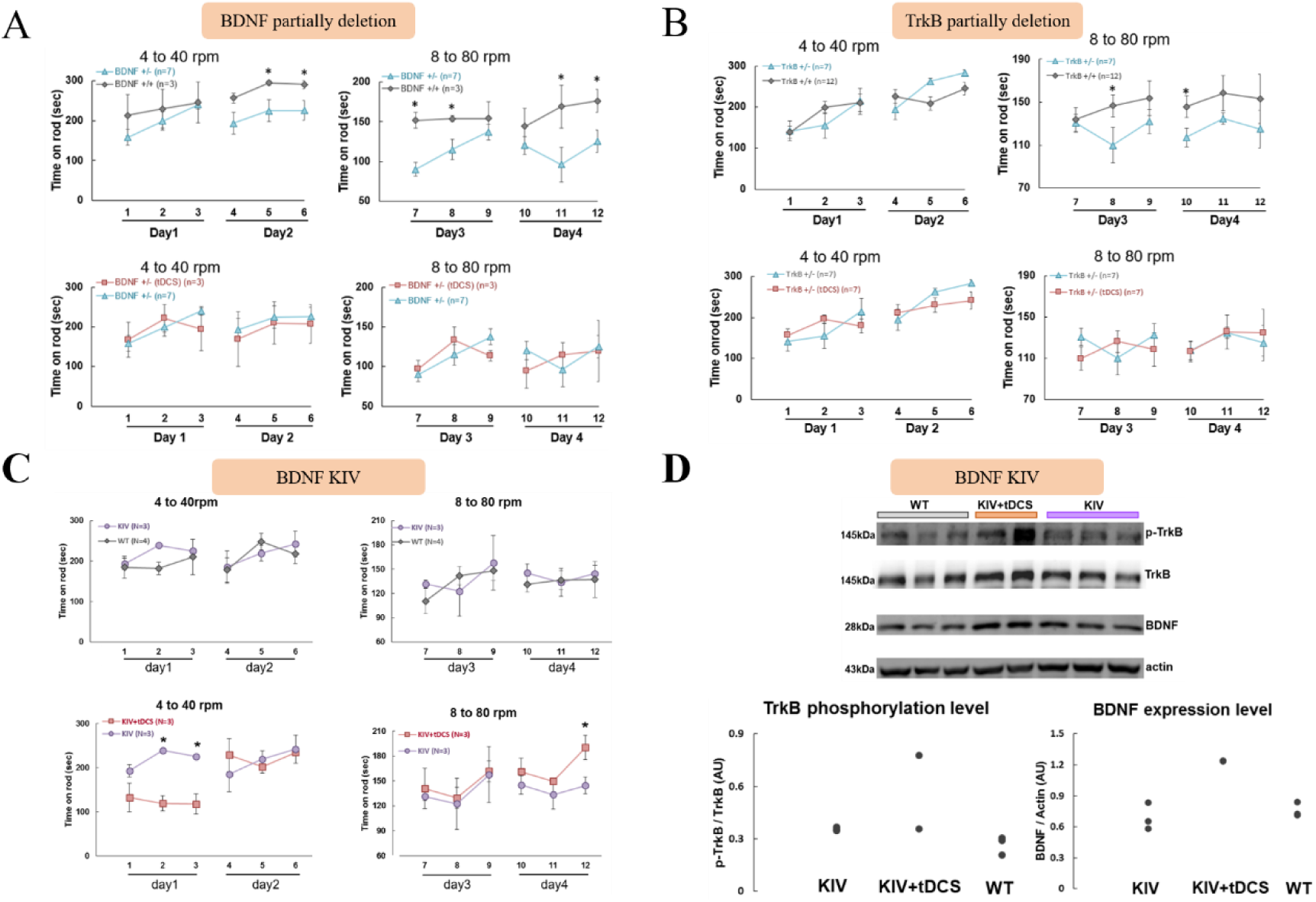
Anodal tDCS modulates rotarod performance in mice with altered BDNF– TrkB signaling. **(A)** Rotarod performance in BDNF heterozygous (BDNF +/−) and wild-type (BDNF +/+) mice without stimulation (upper panels) and during concurrent anodal tDCS (lower panels). **(B)** Rotarod performance in TrkB heterozygous (TrkB +/−) and wild-type (TrkB +/+) mice without stimulation (upper panels) and during concurrent anodal tDCS (lower panels). **(C)** Rotarod performance in BDNF-KIV mice compared with wild-type mice (upper panels), and in BDNF-KIV mice with or without concurrent anodal tDCS (lower panels). For the behavioral experiments in A–C, mice underwent three trials per day over four consecutive days. The rotarod accelerated from 4 to 40 rpm on days 1–2 and from 8 to 80 rpm on days 3–4. Performance was measured as time spent on the rod. **(D)** Representative western blots and quantification of phosphorylated TrkB (p-TrkB), total TrkB, and BDNF in the targeted primary motor cortex (M1) of wild-type mice and BDNF-KIV mice with or without concurrent anodal tDCS. Tissue was collected immediately after completion of the final rotarod training session on day 4; in the tDCS group, stimulation was delivered concurrently with task performance. p-TrkB signals were normalized to total TrkB, and BDNF signals were normalized to actin. Data are presented as mean ± SEM. Sample sizes are indicated in the corresponding panels. Statistical comparisons were performed using independent-samples t-tests. *P < 0.05 for the indicated comparisons. WT, wild type; SEM, standard error of the mean; tDCS, transcranial direct current stimulation.

TrkB heterozygous mice showed numerically lower performance than wild-type mice during several trials at 8–80 rpm, although these differences were not statistically significant. No significant genotype differences were detected in the cohorts receiving concurrent anodal tDCS (Fig. 4B).

BDNF-KIV mice and wild-type mice showed comparable performance without stimulation. In BDNF-KIV mice, the effects of concurrent anodal tDCS varied across trials: stimulated mice performed less well at selected early trials but remained on the rod longer at the final trial on day 4 (Fig. 4C).

Finally, we examined molecular measurements in the targeted M1 of wild-type and BDNF-KIV mice. Tissue was collected immediately after the final training session on day 4, with stimulation delivered concurrently with task performance in the tDCS group. Individual measurements of TrkB phosphorylation and BDNF levels are presented in Fig. 4D. Given the small number of samples and the absence of indicated statistical comparisons, these molecular findings are reported descriptively.

## Discussion

This study examined the behavioral and molecular responses to anodal tDCS delivered at different times relative to motor training. Online stimulation improved rotarod performance at selected trials and increased footstep frequency, whereas stimulation delivered during intertrial intervals did not produce a statistically significant behavioral improvement. Improved performance was also observed at a stimulation-free follow-up two weeks after training. Online stimulation increased TrkB phosphorylation in the stimulated cortex after both one and four days. However, paused offline stimulation also increased TrkB phosphorylation after one day despite lacking a detectable behavioral benefit. Together, these observations suggest that the behavioral effects of stimulation depend on its relationship to motor practice, while increased cortical TrkB phosphorylation alone does not distinguish stimulation schedules associated with improved performance.

The behavioral findings are consistent with the importance of pairing stimulation with ongoing motor practice(*17*). In humans, anodal tDCS delivered during repeated motor training can enhance skill acquisition through changes in between-session consolidation(*2*). Our previous mouse study demonstrated task-specific enhancement of motor learning with concurrent stimulation and preferential facilitation of task-related M1 neurons(*12*). Experimental work in brain slices further showed that direct current stimulation can enhance Hebbian potentiation while preserving pathway specificity and associativity(*9*). These findings provide a rationale for applying stimulation while the relevant circuits are engaged, although they do not establish that online stimulation is superior across all tasks and stimulation configurations.

The stepping analysis extends this behavioral characterization by showing increased footstep frequency alongside improved time on the rod (*18*). Footstep frequency describes movement execution and does not independently establish memory formation. However, improved performance during the stimulation-free assessment on day 18 indicates that the observed benefit was not confined to the period of active stimulation. This finding is consistent with a lasting consequence of training with concurrent tDCS. Because retention was assessed at a single follow-up time point, the present data do not establish the onset, continuity, or maximum duration of this benefit.

One possible explanation for the timing dependence is that direct current interacts with the neural activity generated by the task. Direct current can modulate neuronal compartments, including cell bodies and axon terminals, with effects depending on electric field orientation and the activated pathway(*5*). More recent single-neuron experiments and modeling indicate that modulation of evoked postsynaptic activity can contribute to the effects of direct current on synaptic potentiation(*10*). A multiscale model incorporating realistic neuronal morphologies and human head anatomy further showed that polarization varies across neuronal compartments, cell types, and stimulation configurations(*6*). These studies support an interpretation in which stimulation alters the responsiveness of active circuits rather than producing a uniform change throughout the stimulated cortex.

Presynaptic mechanisms may also contribute. Experimental and modeling work has implicated calcium channels in direct current-induced changes in spontaneous vesicle release(*7*). More recently, measurements in cultured neurons demonstrated that subthreshold electric fields can bidirectionally modulate neurotransmitter release through axonal polarization(*8*). These findings broaden the possible cellular mechanisms through which stimulation could interact with task-related activity. However, they concern preparations distinct from the present awake-mouse experiments and do not demonstrate that altered presynaptic release accounts for our behavioral results. Neural activity, transmitter release, and synaptic potentiation were not measured directly in the cohorts reported here.

The molecular results identify increased TrkB phosphorylation as a detectable response in the stimulated cortex. Increased p-TrkB relative to total TrkB was observed after one day and remained detectable after the fourth day of online stimulation. Because tissue was collected after the final training and stimulation session, these observations demonstrate a response following repeated stimulation, rather than persistence for four days after stimulation cessation. They also do not establish a progressively greater response from day 1 to day 4. BDNF levels were not significantly increased at either sampled endpoint. Thus, the clearest biochemical finding is increased TrkB phosphorylation rather than increased tissue BDNF abundance.

This distinction helps place our results in the context of previous studies. Fritsch et al. (*13*) demonstrated that direct current-induced potentiation in motor cortical slices requires activity-dependent BDNF secretion and TrkB activation. Podda et al. (*14*) reported enhanced hippocampal plasticity and memory alongside epigenetic regulation of Bdnf expression following anodal tDCS. These studies examined different preparations, brain regions, and stimulation schedules. Our cortical protein measurements therefore address a related but distinct question: whether stimulation during motor training is accompanied by changes in tissue BDNF levels and TrkB phosphorylation.

Importantly, tissue BDNF abundance does not directly measure its release or local biological availability. Previous studies have demonstrated that neuronal activity can induce synaptic BDNF secretion and that activity-dependent release differs between axonal and dendritic compartments(*19, 20*). Furthermore, imaging experiments at individual dendritic spines demonstrated an autocrine mechanism in which activity-dependent BDNF release activates TrkB on the stimulated spine and contributes to structural and functional plasticity(*21*). Together, these findings highlight the spatially restricted nature of BDNF signaling and provide a basis for distinguishing localized BDNF–TrkB activity from changes detectable in bulk tissue homogenates. Accordingly, the absence of a significant increase in tissue BDNF in our study does not exclude altered BDNF release or signaling. Nevertheless, increased TrkB phosphorylation alone cannot establish increased BDNF secretion, which was not measured here.

A particularly informative observation is that paused stimulation increased cortical TrkB phosphorylation without significantly improving rotarod performance. This pattern argues against using bulk TrkB phosphorylation alone as a marker of behavioral efficacy. One possibility is that stimulation engages signaling processes that support plasticity, while behavioral improvement additionally requires their temporal overlap with task-related circuit activity. The activity dependence of direct current-enhanced synaptic potentiation provides experimental support for considering this possibility(*9, 10, 13*). Alternatively, online and paused stimulation may differ in the cellular distribution, time course, or downstream consequences of receptor activation. These explanations remain hypotheses; our measurements do not establish equivalent molecular responses between the two protocols.

Preceding online anodal stimulation with brief cathodal stimulation did not provide a detectable additional behavioral benefit. This finding contrasts with a previous human study reporting that cathodal priming before online anodal tDCS enhanced subsequent motor performance and offline learning, although no significant additional benefit was observed during training(*22*). Differences in stimulation duration, training paradigms, and assessment timing may contribute to these divergent findings. Our results do not demonstrate equivalence between the protocols or exclude an effect of other priming intensities, durations, or intervals. More generally, these findings suggest that adding another stimulation phase does not necessarily produce a larger behavioral benefit.

No significant changes in TrkB phosphorylation or BDNF levels were detected in the additional sampled non-targeted regions. This pattern is consistent with a more readily detectable biochemical response near the stimulation site. However, small sample sizes, endpoint sampling, and tissue averaging may limit sensitivity to weaker, transient, or spatially restricted effects. The regional results should therefore be interpreted as the distribution of detectable responses under the conditions examined. Other cellular processes may also contribute to stimulation-induced plasticity: astrocytic calcium signaling has been implicated in the enhancement of cortical responses by tDCS(*15*).

The genetic experiments further explored the relationship between BDNF–TrkB signaling and motor performance. BDNF–TrkB signaling plays an important role in activity-dependent synaptic plasticity and neural circuit function(*23*), while genetic variation affecting activity-dependent BDNF secretion has been associated with altered learning-related neural function and memory performance in humans(*24*). In the present study, BDNF heterozygous mice showed lower rotarod performance than wild-type controls at several trials, while TrkB heterozygous mice showed differences at selected trials. These findings may reflect differences in coordination, initial task proficiency, or acquisition across repeated trials, which the present experiments do not fully separate. Likewise, the absence of statistically significant stimulation effects within a mutant group does not establish that the pathway is necessary for the behavioral benefit. A direct comparison of stimulation responses across genotypes, including a genotype-by-stimulation interaction, would be needed to support a genotype-specific effect.

BDNF-KIV mice showed trial-dependent responses to online stimulation, including lower performance at selected early trials and higher performance at the final trial on day 4. This pattern does not support uniform enhancement throughout training. In the classical BDNF-KIV model, promoter IV-dependent production of BDNF is disrupted, while expression from other promoters remains possible(*25*). Consequently, a behavioral response in this model would not demonstrate that tDCS acts independently of BDNF. Residual BDNF production, developmental adaptations, or altered circuit function could contribute to the observed responses. The limited molecular measurements do not resolve these possibilities.

The cortical measurements provide a complementary physical characterization of the stimulation configuration. The recording-site measurements indicate substantial electrical exposure beneath the epicranial electrode, although the estimated transmission ratio depends on the measured quantity, electrode contact area, tissue preparation, and recording circuit. It should not be interpreted as the fraction of applied current entering the entire brain. Human intracranial measurements demonstrate that transcranial stimulation generates spatially distributed electric fields shaped by anatomy and electrode configuration(*26*). Multiscale modeling further indicates that neuronal polarization depends on both the local field and neuronal morphology(*6*). Our measurements therefore characterize local exposure without providing a complete map of cortical electric fields or their cellular effects.

Several limitations constrain the mechanistic conclusions. Behavioral comparisons at individual trials do not establish an overall change in learning trajectory, and repeated testing requires consideration of multiplicity. Although the stimulation-free follow-up supports a lasting behavioral benefit, additional time points would be needed to characterize retention more fully. Molecular measurements at selected endpoints cannot reconstruct signaling dynamics during stimulation and practice. Bulk tissue assays do not identify responding cell types or distinguish task-engaged circuits from surrounding tissue. Finally, the small and unequal genetic cohorts do not provide definitive tests of pathway necessity or sufficiency. Together, these findings associate concurrent anodal tDCS with improved rotarod performance, increased footstep frequency, a behavioral benefit detectable two weeks after training, and increased TrkB phosphorylation in the stimulated cortex. The molecular response to paused stimulation indicates that pathway engagement and behavioral benefit are related but distinct outcomes. The results support investigating how stimulation timing aligns plasticity-related signaling with task-engaged circuits, while leaving the causal contribution of BDNF–TrkB signaling to be established.

## Acknowledgements

We thank Drs. Chun Xu, Liping Wang, Huatai Xu, and Zhiqi Xiong for valuable suggestions.

## Supplementary figures

**Supplementary Fig. 1.**
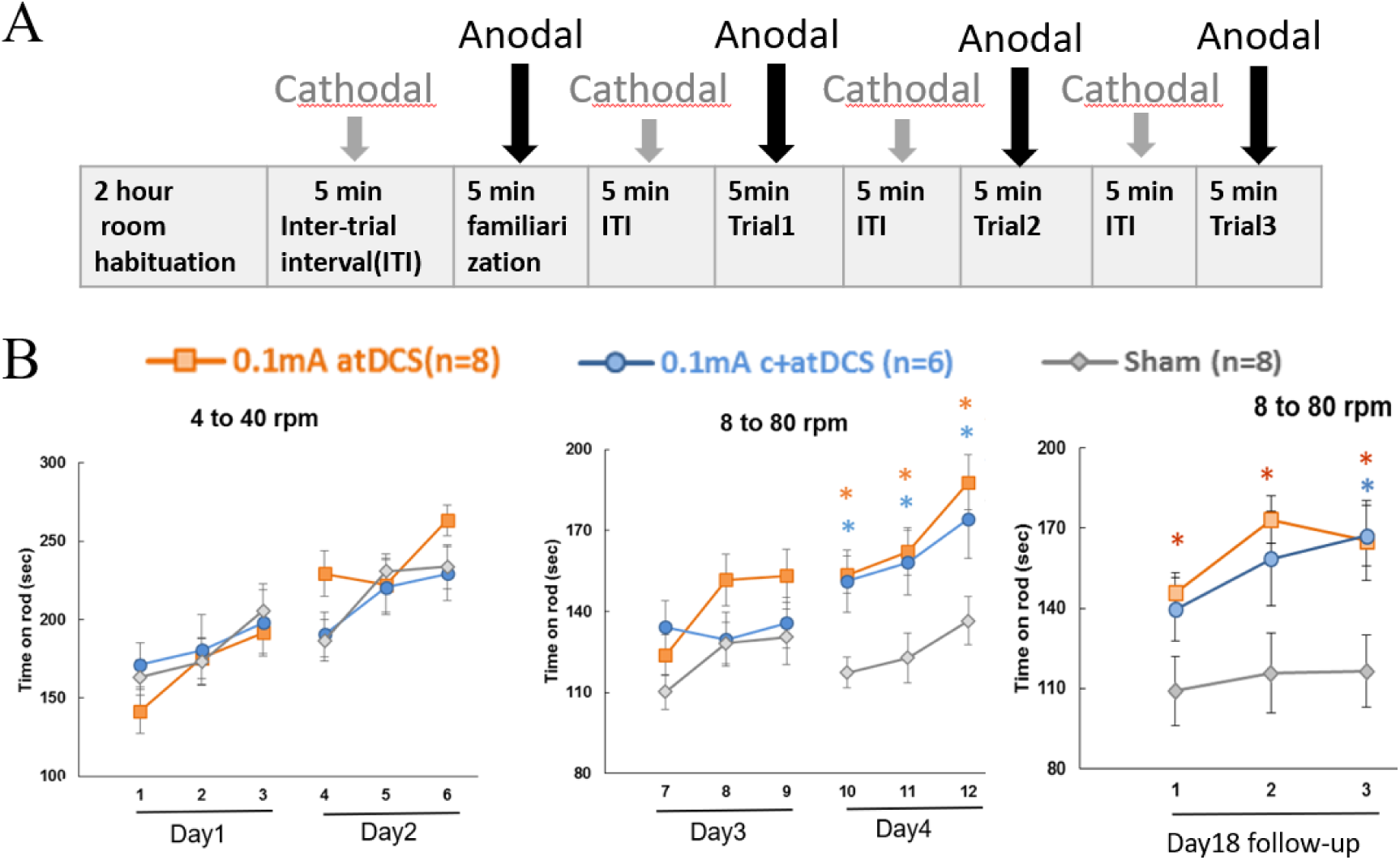
The behavioral benefits of online anodal tDCS remain detectable two weeks after training, with or without preceding cathodal stimulation. **(A)** Experimental timeline for the combined cathodal–anodal stimulation protocol. Following 2 h of room habituation, mice received 5 min of cathodal tDCS before familiarization and before each rotarod training trial. Anodal tDCS was initiated at the start of familiarization and each training trial and continued for 5 min, regardless of early trial termination. Each daily training session consisted of three trials separated by 5-min intervals. **(B)** Rotarod performance in mice receiving online anodal tDCS alone (orange; n = 8), preceding cathodal tDCS followed by online anodal tDCS (blue; n = 6), or sham stimulation (gray; n = 8). Both anodal and cathodal stimulation were delivered at 0.1 mA. The left and middle plots show performance during four consecutive training days, with the rotarod accelerating from 4 to 40 rpm on days 1–2 and from 8 to 80 rpm on days 3–4. The right plot shows performance during three follow-up trials at 8–80 rpm on day 18, 14 days after completion of training and stimulation. No tDCS was delivered during follow-up. Performance was measured as time spent on the rod. Data are presented as mean ± SEM. Statistical comparisons were performed using independent-samples *t*-tests. Orange and blue asterisks indicate significant differences between the respective stimulation groups and the sham group at the corresponding trials. *P* < 0.05. n denotes the number of mice. atDCS, anodal transcranial direct current stimulation; c+atDCS, cathodal stimulation followed by anodal stimulation; ITI, intertrial interval.

**Supplementary Fig. 2.**
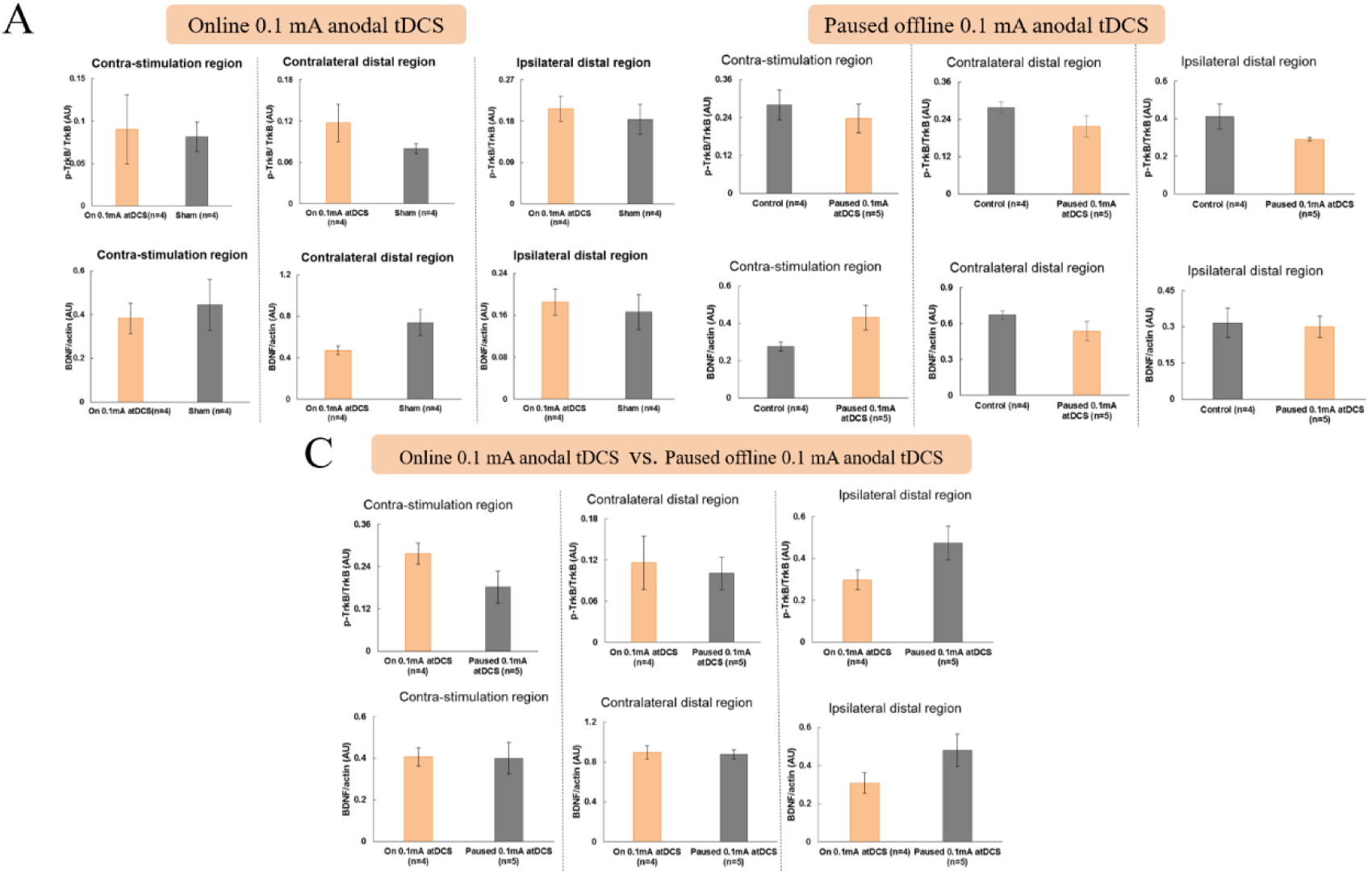
No significant changes in TrkB phosphorylation or BDNF levels are detected in sampled non-targeted brain regions following online or paused anodal tDCS. **(A)** Quantification of phosphorylated TrkB (p-TrkB) relative to total TrkB (upper panels) and BDNF relative to actin (lower panels) in mice receiving paused anodal tDCS (0.1 mA; n = 5) and control mice (n = 4). **(B)** Corresponding measurements in mice receiving online anodal tDCS (0.1 mA; n = 4) and sham stimulation (n = 4). **(C)** Comparisons between online and paused anodal tDCS groups. For each comparison, samples were collected from the region contralateral to the stimulation site, a contralateral distal region, and an ipsilateral distal region, shown from left to right. Online stimulation was delivered concurrently with rotarod task performance, whereas paused stimulation was delivered during intertrial intervals. p-TrkB signals were normalized to total TrkB, and BDNF signals were normalized to actin. Data are presented as mean ± SEM. Statistical comparisons were performed using independent-samples *t*-tests. No statistically significant differences were detected in the comparisons shown. n denotes the number of mice. AU, arbitrary units; tDCS, transcranial direct current stimulation.

## References

1. M. A. Nitsche, W. Paulus, Excitability changes induced in the human motor cortex by weak transcranial direct current stimulation. The Journal of Physiology 527, 633–639 (2000).

2. Janine Reis, Heidi M. Schambra, L. G. Cohen, Noninvasive cortical stimulation enhances motor skill acquisition over multiple days through an effect on consolidation. Proc Natl Acad Sci U S A 106, 1590–1595 (2009).

3. C. Allman*, U. Amadi*, A. M. Winkler, L. Wilkins, N. Filippini, U. Kischka, C. J. Stagg† Heidi Johansen-Berg†, Ipsilesional anodal tDCS enhances the functional benefits of rehabilitation in patients after stroke. Science Translational Medicine 8, 330re331 (2016).

4. G. Schlaug, C. Cassarly, J. A. Feld, S. L. Wolf, V. T. Rowe, S. Fritz, P. Y. Chhatbar, A. Shinde, Z. Su, J. P. Broderick, R. Zorowitz, O. Awosika, D. Edwards, C. Lin, G. E. Franciso, G. F. Wittenberg, S. Pundik, C. Gregory, M. R. Borich, V. Ramakrishnan, W. Feng, Safety and efficacy of transcranial direct current stimulation in addition to constraint-induced movement therapy for post-stroke motor recovery (TRANSPORT2): a phase 2, multicentre, randomised, sham-controlled triple-blind trial. Lancet Neurol 24, 400–412 (2025).

5. A. Rahman, D. Reato, M. Arlotti, F. Gasca, A. Datta, L. C. Parra, M. Bikson, Cellular effects of acute direct current stimulation: somatic and synaptic terminal effects. J Physiol 591, 2563–2578 (2013).

6. A. S. Aberra, R. Wang, W. M. Grill, A. V. Peterchev, Multi-scale model of axonal and dendritic polarization by transcranial direct current stimulation in realistic head geometry. Brain Stimul 16, 1776–1791 (2023).

7. S. O. Vasu, H. Kaphzan, Calcium channels control tDCS-induced spontaneous vesicle release from axon terminals. Brain Stimul 15, 270–282 (2022).

8. A. S. Aberra, M. W. Miles, M. B. Hoppa, Subthreshold electric fields bidirectionally modulate neurotransmitter release through axon polarization. Brain Stimul 19, 103006 (2026).

9. G. Kronberg, A. Rahman, M. Sharma, M. Bikson, L. C. Parra, Direct current stimulation boosts hebbian plasticity in vitro. Brain Stimul 13, 287–301 (2020).

10. F. Farahani, G. Kronberg, M. FallahRad, H. V. Oviedo, L. C. Parra, Effects of direct current stimulation on synaptic plasticity in a single neuron. Brain Stimul 14, 588–597 (2021).

11. David Liebetanz, Michael A. Nitsche, Frithjof Tergau, W. Paulus, Pharmacological approach to the mechanisms of transcranial DC-stimulation-induced after-effects of human motor cortex excitability. Brain 125, 2238–2247 (2002).

12. Y. Wang, J. Wang, Q. F. Zhang, K. W. Xiao, L. Wang, Q. P. Yu, Q. Xie, M. M. Poo, Y. Wen, Neural Mechanism Underlying Task-Specific Enhancement of Motor Learning by Concurrent Transcranial Direct Current Stimulation. Neurosci Bull 39, 69–82 (2023).

13. B. Fritsch, J. Reis, K. Martinowich, H. M. Schambra, Y. Ji, L. G. Cohen, B. Lu, Direct Current Stimulation Promotes BDNF-Dependent Synaptic Plasticity: Potential Implications for Motor Learning. Neuron 66, 198–204 (2010).

14. M. V. Podda, S. Cocco, A. Mastrodonato, S. Fusco, L. Leone, S. A. Barbati, C. Colussi, C. Ripoli, C. Grassi, Anodal transcranial direct current stimulation boosts synaptic plasticity and memory in mice via epigenetic regulation of Bdnf expression. Sci Rep 6, 22180 (2016).

15. H. Monai, M. Ohkura, M. Tanaka, Y. Oe, A. Konno, H. Hirai, K. Mikoshiba, S. Itohara, J. Nakai, Y. Iwai, H. Hirase, Calcium imaging reveals glial involvement in transcranial direct current stimulation-induced plasticity in mouse brain. Nat Commun 7, 11100 (2016).

16. Agnieszka Balkowiec 1, D. M. K. 1, Cellular Mechanisms Regulating Activity-Dependent Release of Native Brain-Derived Neurotrophic Factor from Hippocampal Neurons. J Neurosci. 22, 10399–10407 (2002).

17. Yang Liu, Jieshi He, Ying Wangd, Zhiwei Xud, Qiuxu Liud, Mu-ming Poo, Q. Xie, Pinch restoration in post-stroke hemiplegia rehabilitation: A case report of motor task-synchronized precision repetitive transcranial magnetic stimulation. Brain Stimulation 19, 103101 (2026).

18. P. E. Rothwell, M. V. Fuccillo, S. Maxeiner, S. J. Hayton, O. Gokce, B. K. Lim, S. C. Fowler, R. C. Malenka, T. C. Sudhof, Autism-associated neuroligin-3 mutations commonly impair striatal circuits to boost repetitive behaviors. Cell 158, 198–212 (2014).

19. Matthias Hartmann, R. H. &, V. Lessmann, Synaptic secretion of BDNF after high-frequency stimulation of glutamatergic synapses. The EMBO Journal 20, 5887–5897 (2001).

20. N. Matsuda, H. Lu, Y. Fukata, J. Noritake, H. Gao, S. Mukherjee, T. Nemoto, M. Fukata, M. M. Poo, Differential activity-dependent secretion of brain-derived neurotrophic factor from axon and dendrite. J Neurosci 29, 14185–14198 (2009).

21. S. C. Harward, N. G. Hedrick, C. E. Hall, P. Parra-Bueno, T. A. Milner, E. Pan, T. Laviv, B. L. Hempstead, R. Yasuda, J. O. McNamara, Autocrine BDNF-TrkB signalling within a single dendritic spine. Nature 538, 99–103 (2016).

22. M. Christova, D. Rafolt, E. Gallasch, Cumulative effects of anodal and priming cathodal tDCS on pegboard test performance and motor cortical excitability. Behav Brain Res 287, 27–33 (2015).

23. H. Park, M.-m. Poo, Neurotrophin regulation of neural circuit development and function. Nature Reviews Neuroscience 14, 7–23 (2012).

24. Michael F. Egan 1, Masami Kojima 7 4 5 6, Joseph H. Callicott 7 1, Terry E. Goldberg 7 1, Bhaskar S. Kolachana 1, Alessandro Bertolino 1, Eugene Zaitsev 4, Bert Gold 3, David Goldman 2, Michael Dean 3, Bai Lu * 4, D. R. W. 1, The BDNF val66met Polymorphism Affects Activity-Dependent Secretion of BDNF and Human Memory and Hippocampal Function. Cell 112, 257–269 (2003).

25. Kazuko Sakata, Newton H. Woo, Keri Martinowich, B. Lu, Critical role of promoter IV-driven BDNF transcription in GABAergic transmission and synaptic plasticity in the prefrontal cortex. Proc Natl Acad Sci U S A 106, 5942–5947 (2009).

26. Y. Huang, A. A. Liu, B. Lafon, D. Friedman, M. Dayan, X. Wang, M. Bikson, W. K. Doyle, O. Devinsky, L. C. Parra, Measurements and models of electric fields in the in vivo human brain during transcranial electric stimulation. Elife 6, (2017).

